# How do King Cobras move across a major highway? Unintentional wildlife crossing structures may facilitate movement

**DOI:** 10.1101/2021.07.30.454480

**Authors:** Max Dolton Jones, Benjamin Michael Marshall, Samantha Nicole Smith, Matt Crane, Inês Silva, Taksin Artchawakom, Pongthep Suwanwaree, Surachit Waengsothorn, Wolfgang Wüster, Matt Goode, Colin Thomas Strine

**Affiliations:** School of Biology, Suranaree University of Technology, Nakhon Ratchasima, Thailand; Thailand Institute of Science and Technological Research, Nakhon Ratchasima, Thailand; School of Bioresources and Technology, King Mongkut’s University of Technology Thonburi, Bangkok, Thailand; Molecular Ecology and Fisheries Genetics Laboratory, School of Natural Sciences, Bangor University, Bangor, UK; School of Natural Resources and Environment, University of Arizona, Tucson, Arizona, USA

**Keywords:** bridge, drainage culvert, mortality, *Ophiophagus hannah*, road crossing, space use

## Abstract

1. Global road networks continue to expand, and the wildlife responses to these landscape-level changes need to be understood to advise long-term management decisions. Roads have high mortality risk to snakes because snakes typically move slowly and can be intentionally targeted by drivers.
2. We investigated how radio-tracked King Cobras (*Ophiophagus hannah*) traverse a major highway in northeast Thailand, and if reproductive cycles were associated with road hazards.
3. We surveyed a 15.3km stretch of Highway 304 to determine if there were any locations where snakes, and other wildlife, could safely move across the road (e.g., culverts, bridges). We used recurse analysis to detect possible road-crossing events, and used subsets of King Cobra movement data to create dynamic Brownian Bridge Movement Models (dBBMM) in an attempt to show movement pathways association with possible unintentional crossing structures. We further used Integrated Step Selection Functions (ISSF) to assess seasonal differences in avoidance of major roads for adult King Cobras in relation to reproductive state.
4. We discovered 32 unintentional wildlife crossing locations capable of facilitating King Cobra movement across the highway. Our dBBMMs failed to show if underpasses were being used by telemetered individuals; however, the tracking locations pre- and post-crossing provided strong evidence of underpass use. Our ISSF suggested a lower avoidance of roads during the breeding season, though the results were inconclusive. With the high volume of traffic, large size of King Cobras and a 98.8% success rate of crossing the road in our study, we strongly suspect that individuals are using the unintentional crossing structures to safely traverse the road.
5. Further research is needed to determine the extent of wildlife underpass use at our study site and globally, alongside using previously proven fencing to facilitate their use. We propose that more consistent integration of drainage culverts and bridges could help mitigate the impacts of roads on some terrestrial wildlife, particularly in areas where roads fragment forests and wildlife corridors.

## Introduction

Southeast Asia is one of the world’s many biodiversity hotspots, combining a rich fauna and flora with a myriad of human-mediated threats are endangering the biodiversity (Myers et al. 2000; Hughes 2017; Ng et al. 2020). The growing human population in Southeast Asia continues to increase urbanisation (Schneider et al. 2015), and road networks continue to expand to meet human demands, posing threats to wildlife (Ascensão et al. 2018; Hughes 2018). Roads are either diffuse or hard barriers to wildlife movement (Shepard et al. 2008a; Brehme et al. 2013), dividing habitats and resources, and potentially undermining wildlife population integrity and reducing genetic diversity (Aresco 2005; Row et al. 2007; Balkenhol and Waits 2009; Clark et al. 2010; Jackson and Fahrig 2011; Herrmann et al. 2017). Alongside fragmenting habitats, roads can also constitute a direct source of mortality for wildlife via vehicular collisions (Bernardino and Dalrymple 1992; Rosen and Lowe 1994; Lodé 2000; Aresco 2005; Das et al. 2007; Row et al. 2007). Some drivers can intentionally target certain animals, such as snakes; therefore, potentially leading to targeted species disproportionately affected by roads (Langley et al. 1989; Ashley et al. 2007; Beckmann and Shine 2012; Assis et al. 2020; but not detected by Secco et al. 2014).

Wildlife managers can alleviate the risks from roads, mitigate road mortality and facilitate animal movement by implementing wildlife-crossing infrastructure (Lister et al. 2015). Once wildlife becomes acclimated to crossing structures, the structures help sustain animal mobility across fragmented landscapes, aiding wildlife access to resources and conspecifics for gene flow (Clevenger and Barrueto 2014). Wildlife-crossing locations can be underpasses, such as culverts or tunnels, or overpasses, which are generally large vegetated land bridges (Dodd Jr. et al. 2004; Clevenger and Huijser 2009; Glista et al. 2009). Wildlife crossings have been successful in facilitating animal movement in several cases (Forman 2003; Beckmann et al. 2010), such as for mule deer in western North America (Simpson et al. 2016), a diversity of wild mammals in Poland (Myslajek et al. 2016, Ważna et al. 2020) and turtle species in Canada (Markle et al. 2017).

Snakes are common victims of roads, likely due to low movement speeds, unique body shape and mode of locomotion (Andrews and Gibbons 2005). Additionally, snakes are disproportionately targeted by road users (Ashley et al. 2007; Beckmann and Shine 2012), and thus an important group to protect from the risks presented by roads. However, studies so far reveal an infrequent, and unpredictable, use of ecopassages by snakes (Baxter-Gilbert et al. 2015).

The King Cobra (*Ophiophagus hannah* [CANTOR, 1836]), is a large, venomous snake widely distributed throughout Southeast Asia and ranging from India to China and the Philippines. The IUCN classifies King Cobras as Vulnerable (Stuart et al. 2012) with decreasing populations and urges investigations into King Cobra’s threats. Andrews and Gibbons (2005) showed that stout-bodied species (in the South eastern USA) had slower crossing speeds than longer, slender-bodied sympatric species. This suggests that King Cobras could also exhibit relatively fast movement speeds across roads; however, these crossing speeds would likely be undermined by the large length and mass of this active forager. Actively foraging species, with high mobility, demonstrate plasticity in their use of microhabitats, often increasing their risk from roads (Forman et al. 2003; Hartmann et al. 2011). King Cobras are susceptible to road mortality in areas where major roads divide habitats, such as in the Sakaerat Biosphere Reserve (SBR) in northeast Thailand (Marshall et al. 2018; 2020). Despite small sample sizes of mortalities in Marshall et al. (2018), four vehicle collisions were recorded among a total of 14 mortality events, prompting further investigation into the potential impacts roads have on King Cobras.

During routine road construction, plans typically integrate drainage culverts sporadically to divert water from road surfaces. However, such structures may also act as unintended wildlife-crossing locations for small taxa (Clevenger and Waltho 2000; Ng et al. 2004; Aresco 2005; Ascensão and Mira 2007; Grilo et al. 2008; Sparks and Gates 2017; Brunen et al. 2020). In central Ontario, Canada Baxter-Gilbert et al (2015), found that three reptile species used culverts as eco-passages during monitoring: Painted Turtles (*Chrysemys picta*), Snapping Turtles (*Chelydra serpentina*) and Northern Watersnakes (*Nerodia sipedon*). In addition, Aresco (2005) demonstrated the importance of an under-highway culvert for reducing turtle mortality, when augmented with drift fences.

Based on the evidence presented above, it is reasonable to suggest that King Cobras are likely using some unintended wildlife crossing structures to safely traverse the roads. The abundance and importance of roads in and around the BR make this an ideal site to explore the role of these structures in assisting movement across roads by King Cobras. In this paper, we therefore aim to identify potential areas that could facilitate movement of these large snakes across a busy major road within the SBR. Using a long-term dataset on the movement ecology of King Cobras in northeast Thailand, we explore the following: 1) are there any structures present along the highway which could facilitate King Cobra movement? 2) are King Cobra reproductive cycles associated with road hazards (e.g., seasonal avoidance of roads or increased rates of vehicle collision)?

## Methods

### Study site

We conducted field work from 22 March 2014 to 28 August 2020, at the Sakaerat Biosphere Reserve (SBR), Nakhon Ratchasima Province, Thailand (14.44-14.55° N, 101.88-101.95° E; Figure 1). The SBR consists of three areas of differing levels of protection: a core protected forested area covering 80 km^2^ consisting of mainly dry dipterocarp forest and dry evergreen forest; a buffer zone consisting of areas of reforestation and plantations, and lastly a transitional area dominated by agriculture (rice, casava, corn, and sugar). The transitional area contains 159 settlements with 72,000 inhabitants (Thailand Institute of Scientific and Technological Research, 2018), and a network of both paved and dirt roads. The forested areas of the core area and transitional zone are bisected by the National Highway 304 (with dense, expansive forest on either side of the highway), built initially in 1956, with further road improvement in 1966 and subsequent expansion from two to four lanes in 2005 (Laurence 2014; Vaeokhaw et al. 2020). Highway 304 transects several protected forests, including the largest fragment of surviving forest in Central Thailand that hosts high biodiversity of threatened or endangered herpetofauna (Silva et al. 2020).

**Figure 1.**
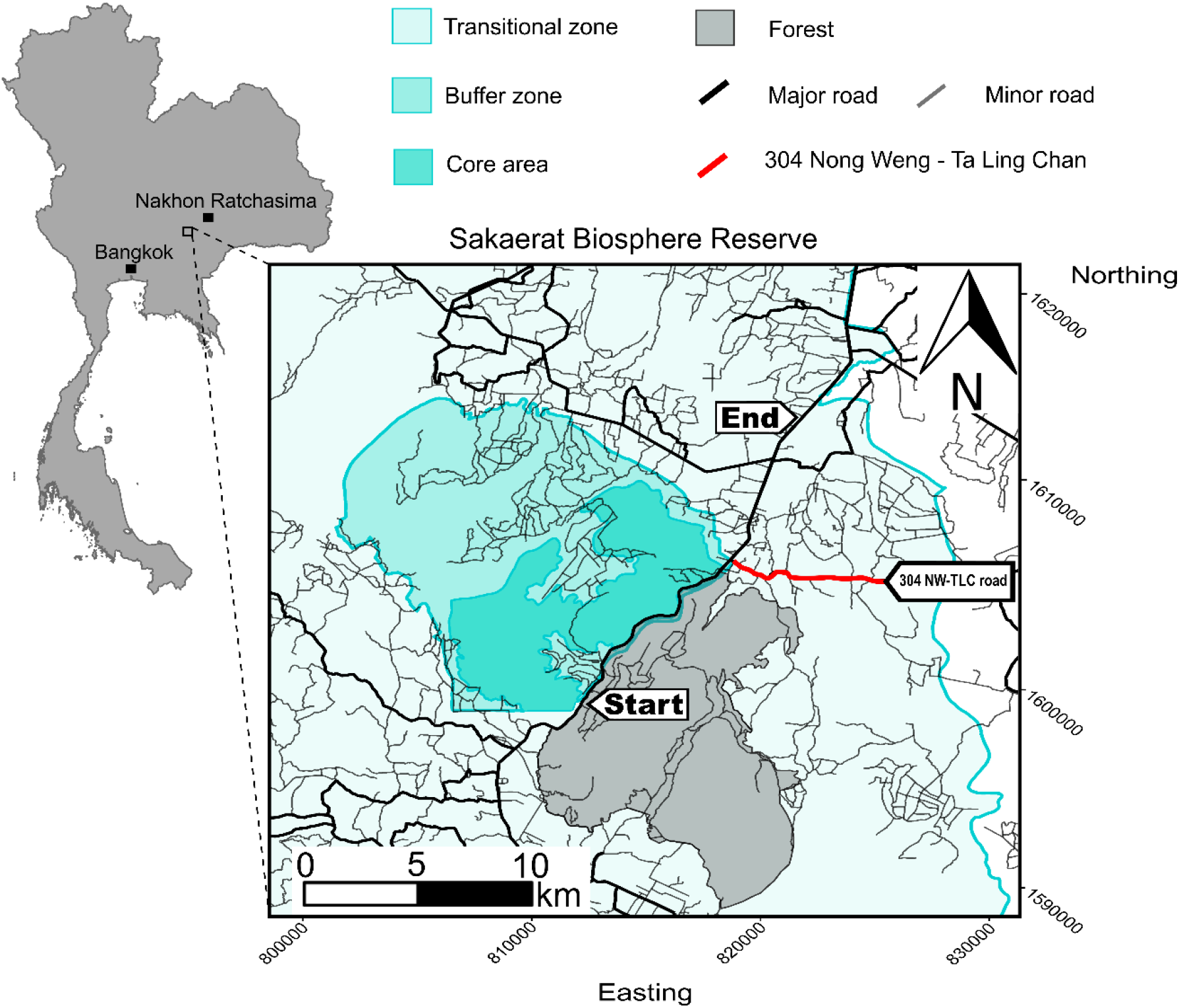
Study site map in relation to Bangkok and Nakhon Ratchasima cities. The three Sakaerat Biosphere Reserve zones are delineated by their level of protection via increased opacity (increasing opacity represents increased protection). The bold red line shows the 304 Nong Weng – Ta Ling Chan Road. The *Start* and *End* mark the section of Highway 304 included in our study.

A second major road perpendicular to the Highway 304, Nong Weng – Ta Ling Chan (304NW-TLC), passes through a populated area in the transitional zone, East of the Highway 304 (Figure 1). We studied this road because it is the first major road (with tarmac and multiple lanes) separating the agricultural area from the un-protected forest fragment to the south; therefore, the road has substantial conservation implications, especially for female King Cobras that are forced to circumvent the road to reach forested areas, where oviposition typically occurs.

### Capture

We performed unstandardized visual encounter surveys, on foot and via road-cruising surveys using motorcycles, throughout accessible locations in the SBR (Marshall et al. 2018, 2019, 2020). We also relied on notifications from local residents and rescue teams to locate and capture individuals. We captured King Cobras between 19 March 2014-18 March 2020, using a combination of opportunistic captures, villager notations and active visual surveys. We gave each snake a unique ID based on age-class, sex and capture number (e.g., AM018 refers to an adult male that was the 18^th^ King Cobra captured, JF055 refers to a juvenile female that was the 55^th^ King Cobra captured).

We anaesthetised King Cobras using isofluorane to obtain accurate morphological measurements and perform radio transmitter implant surgeries, following procedures outlined in Reinert and Cundall (1982). We implanted Holohil AI-2T or SI-2T transmitters into the coelomic cavity. We initially marked individuals with a unique brand on the ventral/dorsal scales using a disposable medical cautery device (Winne et al. 2006). We switched to passive integrated transponders (PIT-tags) beginning with AM054.

We released snakes within 24 hours post-surgery, at an average distance of 191 m (range = 0 – 1263 m, Supplementary Table 1) from their capture site. We aimed to release snakes as close to their reported capture site as possible; however, distances had to be increased when snakes were captured within, or near homes. Our largest recorded distance released from capture location was the result of needing to remove the snake from a large human settlement area, as requested by residents. All release locations were subsequently shown to be within the estimated occurrence distributions of our telemetered individuals. Because we recaptured AM006, AM007 and AF010 after transmitters from their first implant failed, we used capture and release information from this subsequent recapture, because the original data were inaccurate (Supplementary Table 1).

### Radio telemetry

Radio tracking protocol changed throughout the study, due to staff availability and changes in investigation targets (initially we only aimed to assess home range sizes and habitat use, while in later years we added movement patterns and site fidelity to the program). We tracked snakes nearly continuously, until AM006. We maintained signal contact with the snakes every 15 minutes, and determined the snake’s location every hour between 06:00-22:00. From 09 March 2014-28 July 2018, we radio tracked individuals 006-026 four times per day (i.e., 06:30, 11:00, 16:00, 20:00), with a mean of 8.5 ± 0.1 hours between fixes. We radio tracked individuals 027-099, from 28 July 2018-01 August 2020 three times per day aiming for 5 h intervals between successful pinpoints (Supplementary Figure 1 displays overall study time lag distribution for each individual). We usually radio tracked King Cobras in daylight; however, we occasionally radio tracked snakes at night, depending on individual movements and landscape type. We triangulated snake locations, attempting to maintain a minimum distance of 10 m from the snake (occasionally compromised by sub 10m GPS accuracy or difficult terrain), which enabled us to be reasonably confident that the snake was within a 5 m^2^ area. We recorded the triangulated location (Universal Transverse Mercator 47 N WGS 84 datum) using handheld GPS units, recording date, time and GPS accuracy.

### Quantifying crossing structure characteristics

We identified drainage culvert locations along Highway 304 using roadside markers, presumably set by construction workers. No over-the-road structures exist at our study area, and therefore all crossing structures mentioned herein refer to corridors which allow animals to move directly underneath the road. We recorded locations of both entrances for all drainage culverts and bridges (viaducts) encountered, along with vertical diameter of entrance (mm), horizontal diameter of entrance (mm), length of structure (m), vegetation cover at entrance (yes/no), dominant substrate within the structure and connectivity to landscape feature (i.e., none, stream or irrigation canal) for each crossing. We calculated distances (m) between adjacent potential crossing structures with the measuring tool in QGIS (v. 3.14.15 ‘pi’).

### Identifying road crossing events

We manually created spatial polygons using QGIS v.3.14.15 ‘pi’, for the entire study area encompassing the side of Highway 304 that contained the core protected area, herein referred to as *North Side* (Supplementary Figure 2). We used the *recurse* package v.1.1.0 in R (Bracis et al. 2018) to calculate the number of times each telemetered snake entered, or exited, the North Side spatial polygon, which corresponded to a road-crossing event across Highway 304 (Supplementary Figure 2). We also created a spatial polygon encompassing the area South of the 304NW-TLC road; herein referred to as *South Side* (Supplementary Figure 3). Using the *recurse* package, we recorded each time a nesting female King Cobra entered, or exited, the South Side spatial polygon, corresponding to an event during which the snake traversed the road (Supplementary Figure 3). Due to only having one adult male which interacted with the 304NW-TLC road and poor temporal resolution for this individual, we chose to only sample adult female King Cobras for the *South Side* spatial polygon. This allowed us to investigate if there were any temporal patterns for reproductive females to interact with the 304NW-TLC road during nesting movements.

The recurse analysis provided the approximate time that a highway crossing event occurred for each snake (although the time provided is restricted to the nearest datapoint collected). We then took subsets of the radio tracking data, that consisted of fixes taken two weeks prior to, and two weeks after, each crossing event. We ran dynamic Brownian Bridge Movement Models (dBBMMs) on these subsets to estimate an occurrence distribution describing the possible movement pathways taken during a crossing event, using the *move* package v.3.1.0 in R (Kranstauber et al. 2016). Because the subsets were shorter time periods than our overall tracking periods, we used a window size of 15 and margin size of 3 to detect temporally fine-scale changes in movement states (specifically, shifts between resting/sheltering and movement) when using underpasses. Following the methods outlined in Marshall et al. (2020), we extracted 90%, 95% and 99% contours (confidence areas), using R packages *adehabitatHR* v.0.4.16 (Calenge 2006), and *rgeos* v.0.4.2 (Bivand and Rundel 2020), to visualise the movement pathways when crossing Highway 304. Examples of our dBBMM subsets can be seen in Supplementary Figure 4-9. In addition to our dBBMM subsets, we also visualised the points directly before and after a crossing event, to investigate if single, direct movements could also serve as a proxy to determine underpass use by adult King Cobras.

### Integrated step-selection function

We used Integrated Step-Selection functions (ISSF) from the *amt* package v.0.0.6 in R (Signer et al. 2019) to assess the influence of major roads on adult King Cobra movement (i.e., avoidance or attraction). Integrated Step-Selection functions use observed locations (steps) of telemetered animals to generate random steps using observed step characteristics (step length, turning angle). In ISSFs, predictor covariates (i.e., Euclidean distance to major roads) are used to discern which covariates influence animal movement.

We separated tracking periods in to two seasons, incorporating breeding and nesting in one season and the remainder of an individual’s tracking duration during the non-breeding season. The earliest date we observed breeding (over multiple years) was 10 March, and the latest date that a female left her nest was 05 July, which we used to define the extent of breeding season. We added 10-day buffers to start and end dates to account for natural variation that we may have missed in the population, ultimately resulting in an annual breeding season from 01 March to 15 July. We used an inverted raster layer which described varying distances from major roads within the SBR; we inverted the raster to aid interpreting model outputs (Marshall et al. 2020). Following methodology by Fortin et al. (2005), Marshall et al. (2020) and Smith et al. (2021), we simulated 200 random points for each step, allowing for broad sampling of the surrounding landscape. We opted to use a large number of random points due to the coarseness of VHF radiotelemetry compared to GPS telemetry data, the latter usually only affording a single, or very few, random steps per used step due to the high temporal resolution of data and computational cost (Northrup et al. 2013; Thurfjell et al. 2014). Two-hundred random points also facilitates coverage of rare features or smaller changes within a landscape.

We evaluated avoidance of, and attraction to, roads by telemetered adult King Cobras at both the individual and population levels. Each model included step length, turning angle and (inverted) distance from major roads as predictors. We investigated population-level effects following R script by Muff et al. (2020), which involved using a Poisson regression model with stratum-specific effects, and accounting for the data’s structure (both individual ID and step/strata) using Gaussian processes. Following Muff et al., (2020) we used fixed prior precision of 0.0001 for strata-specific effects and fitted the model using the *INLA* v.20.03.17 package (Rue et al. 2020). We radio tracked AF010, AF096 and AF099 only during the breeding season, so these snakes were only included in the breeding models. In contrast, we radio tracked AF056 only during the non-breeding season and therefore was only used in non-breeding season models. We included AF017, AF058 and AF086 in both non-breeding and breeding models. In summary, we included four adult females in population-level non-breeding models and six adult females in population-level breeding models. All adult males had sufficient data to be included in all ISSF models.

### Software

Information about the software and R packages used can be found in the supplementary document. We opted to follow best practises in data sharing to enhance utility of our study and optimise for meta-analysis inclusion (Tedersoo et al. 2021).

## Results

### Radiotelemetry

From 22 March 2014-28 July 2020, we radio tracked 21 King Cobras: eight adult males, seven adult females, four juvenile males, and two juvenile females (Supplementary Table 2). We recaptured, and subsequently radio tracked, three snakes (i.e., AM006, AM007 and AM010), after 842, 1405, and 280 days missing from the study respectively. We radio tracked snakes for an average of 344.53 ± 55.65 days (range = 134 – 3122 days). We obtained an average of 920 ± 157 fixes (range = 66 – 1176 fixes) on telemetered King Cobras, with an average of 9 ± 0.06 hours (range = 0.05 – 793.85 hours; Supplementary Figure 1) between fixes. Snakes relocated on average 263 ± 48 times during telemetry (range = 31 – 985 relocations).

### Road-crossing location and characteristics

We recorded 32 potential road-crossing locations (underpasses) along the 15.3 km section of highway (Euclidean distance from the first and last crossing point; Figure 2). Of the 32 road-crossing locations, 21 were single drainage culverts, seven were double drainage culverts (two culverts side by side) and four were bridges (Figure 2). Twenty-six of these crossing points (24 drainage culverts and two bridges) were within an 8 km section of the highway adjacent to the forest comprising the protected area of the SBR (Figure 2). Road-crossing structures were spaced out along the highway at a mean distance of 536.3 ± 88.4 m (range = 191 – 2620 m).

**Figure 2.**
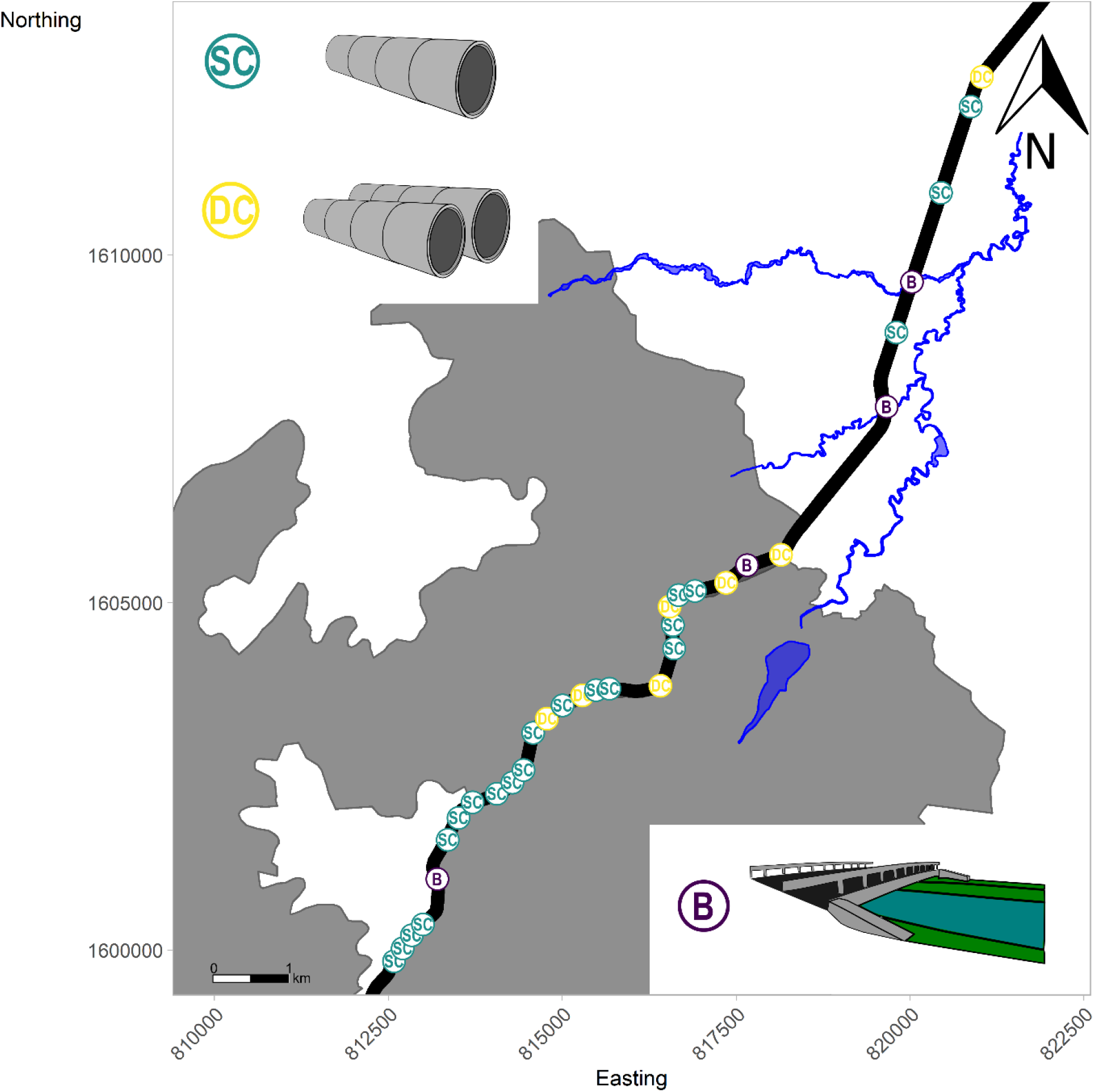
Location and structure type of all crossing structures throughout the survey area. *SC* = single culvert, *DC* = double culvert, *B* = bridge. Grey depicts forested areas and blue indicates irrigation canals and water features throughout the site. Culverts are named in chronological order (C1-C32) from southwest to northeast (Supplementary Table 2).

Crossing structures had a mean length of 40.94 ± 1.75 m (range = 26 – 82 m), a mean entrance height of 1138.16 ± 127.13 mm (range = 194 – 3000 mm), and a mean entrance width of 3792.22 ± 1418.51 mm (range = 543 – 30000 mm; Supplementary Table 2). All crossing points were concrete constructions, except for one metal drainage culvert (C24). There was usually no substrate within structures (n = 17), although those with substrate consisted of gravel (n = 5), rocks (n = 4), water (n = 3), soil (n = 2) and in one instance tar-like substance. Most crossing structures were not connected to any further water flow systems (n = 20); nine were adjacent to stream beds and three had connecting irrigation canals (three out of the four bridges were connected to irrigation canals). All crossing structures contained some anthropogenic waste either at the entrance, or within the structure. The entrances of only four culverts were devoid of any vegetation cover (C4, C6, C14 and C28).

### Road Crossing and motion variance

We confirmed nine of 21 telemetered King Cobras had crossed Highway 304: five adult males (AM006, AM007, AM015, AM018, AM054), three adult females (AF010, AF017, AF058) and one juvenile female (JF055; Figure 3). Adult males crossed the highway 14 times (range = 1 – 37 times per individual) on average, adult females crossed an average of twice (range = 2 – 3 times per individual) and the single juvenile female crossed four times. We ultimately recorded 84 crossing attempts of Highway 304, with one King Cobra fatality; thus, resulting in a 98.8% success rate when attempting to traverse the road (Figure 4).

**Figure 3.**
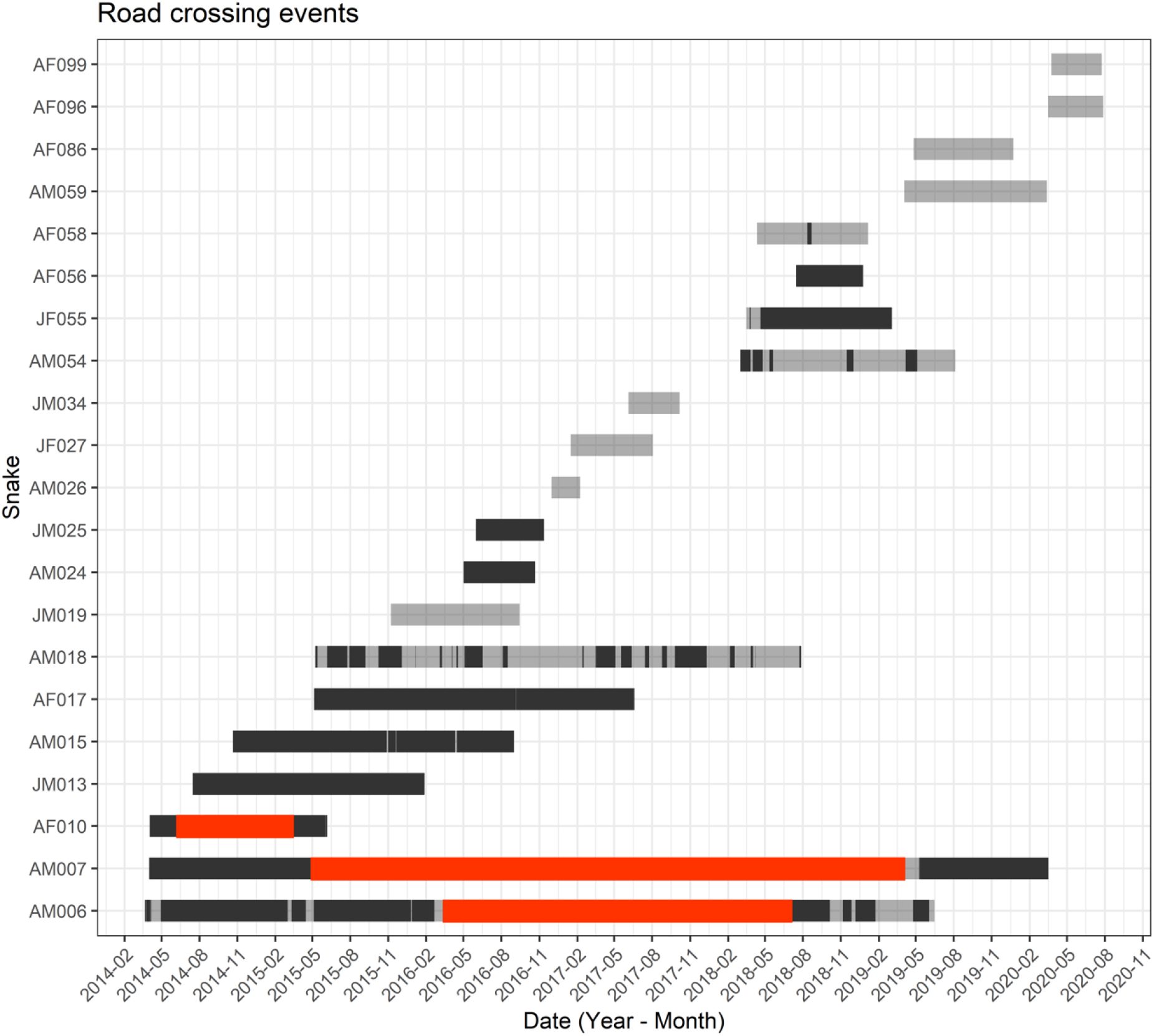
Road-crossing events from all 21 telemetered King Cobra. Each grey bar corresponds to an individual, and opaque bars show when individuals were within the North Side spatial polygon. Transitions from grey to black correspond to a snake crossing over Highway 304. Red bars indicate periods of time when individuals were not radio tracked.

**Figure 4.**
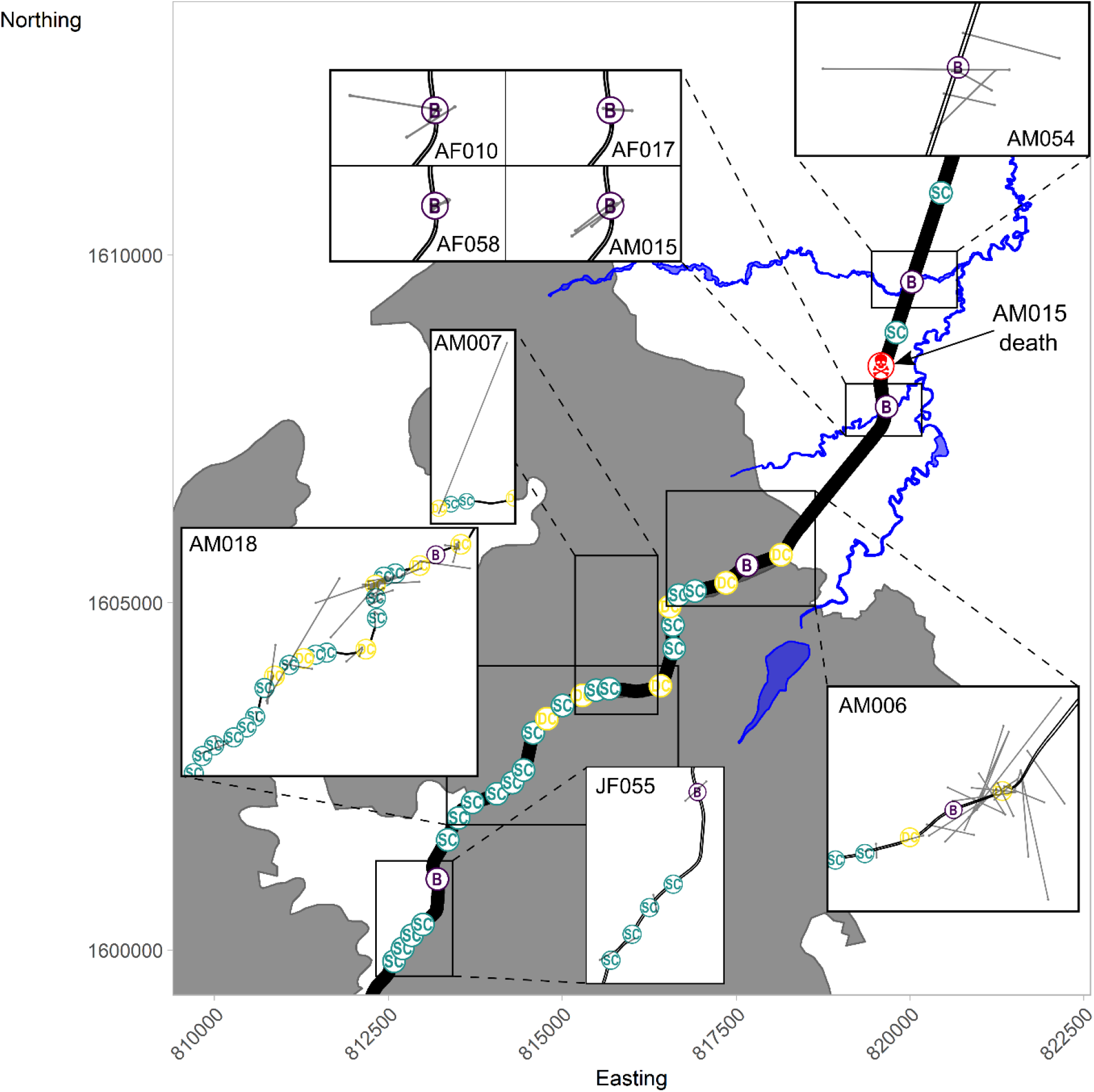
All recorded King Cobra road-crossings along Highway 304, overlapped with crossing structure locations and structure type. *SC* = single culvert, *DC* = double culvert, *B* = bridge. The map also depicts the location of AM015’s death (road-kill).

Each telemetered female that nested on the South side of the highway, crossed over 304NW-TLC at least once during the study. All nesting females South of the road entered the South Side spatial polygon, and associated forested/forest-adjacent area, between 11 April-5 May. Three of four telemetered females subsequently left the South Side spatial polygon, which corresponds to moving away from forested area and into the agricultural matrix, from 18 June-2 July. One female, AF099, moved North following successful nesting, making continuous movements towards 304NW-TLC road; however, her transmitter failed on 24 July 2020 before we could observe her crossing the road and leaving the forest. We radio tracked three females, AF058, AF086 and AF096, for 182, 188 and 40 days respectively after they crossed back to the North side of 304NW-TLC road. During this subsequent radio tracking, females used the agricultural landscape and we recorded no further crossing events (Figure 5).

**Figure 5.**
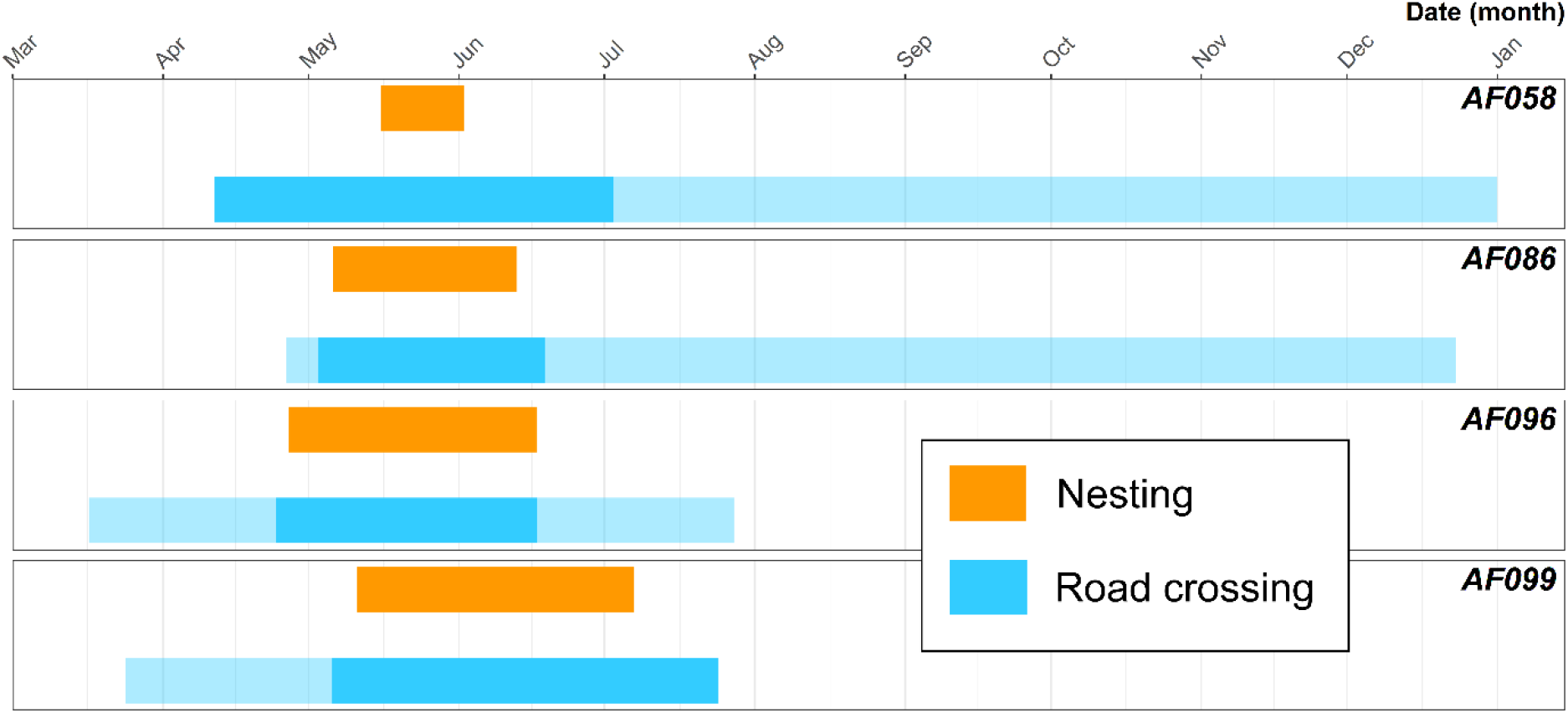
Adult female King Cobra nesting and road crossing. Orange bars highlight the nesting duration. Opaque blue bars represent when an individual was within the South Side spatial polygon (South of 304NW-TLC road within forested areas for nesting). Transitions from translucent to opaque blue bars show road crossing events.

### Integrated step-selection functions

Because we inverted our raster layers (Euclidean distances to major roads), positive coefficients expressed positive association with roads (i.e., attraction). Locations of nine adult King Cobras were positively associated with major roads during the breeding season (AF010, AF017, AF058, AF096, AM006, AM007, AM018, AM024 and AM059; Figure 6); whereas, locations of four individuals indicated avoidance of roads during the breeding season (AF086, AF099, AM015 and AM054; Figure 6). Four adult males showed a positive association with roads during the non-breeding season (AM007, AM015, AM018 and AM024; Figure 6). Three adult males and all four females included in the non-breeding season model exhibited an avoidance of major roads (AF017, AF056, AF058, AF086, AM006, AM054 and AM059; Figure 6). However, all confidence intervals for adult females overlapped zero, and many of our results for the adult males, likely due to our coarse radiotelemetry fixes which limited our inferences.

**Figure 6.**
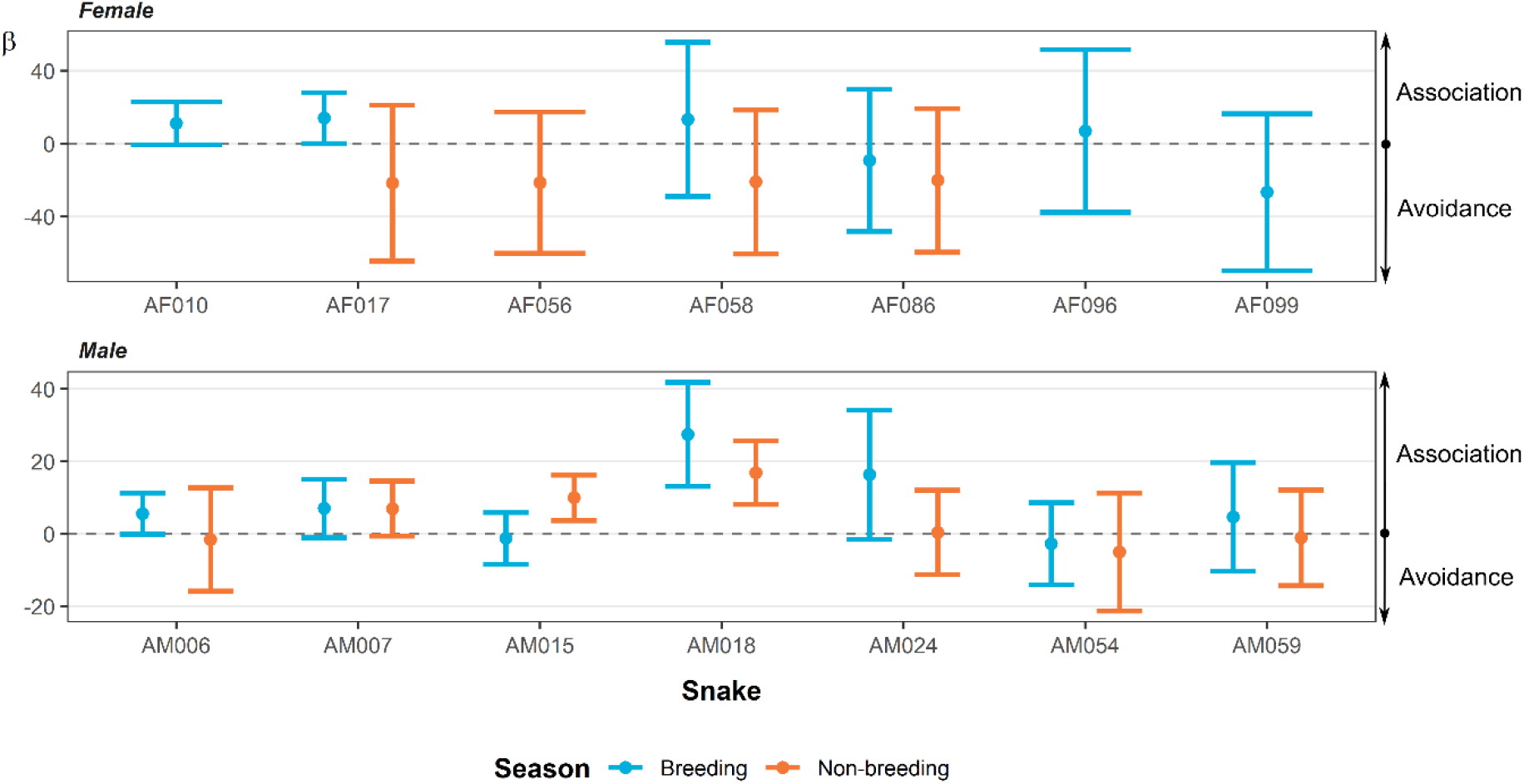
The coefficients relating to major roads from the integrated step-selection function analyses. Breeding and non-breeding season are depicted by blue and orange respectively. Circles show the β estimate from the model and error bars show the associated 95% confidence intervals for each estimate.

Population-level ISSF models indicated low association with major roads during the breeding season in adult males (β = 4.76^-04^, 95% CI -5.42^-04^ – 0.0014; Figure 7) and adult females (β = 5.02^-04^, 95% CI -1.52^-04^ – 0.0012; Figure 6). The population-level ISSF for adult males in the non-breeding season showed a similar result to the breeding season (β = 4.83^-04^, 95% CI -1.05^-04^ – 0.001; Figure 7); however, the adult females exhibited a lower association during the non-breeding season (β = 1.9^-04^, 95% CI -9.61^-04^ – 0.0013; Figure 7). In all population-level ISSF models, confidence intervals overlapped zero which limits inferences.

**Figure 7.**
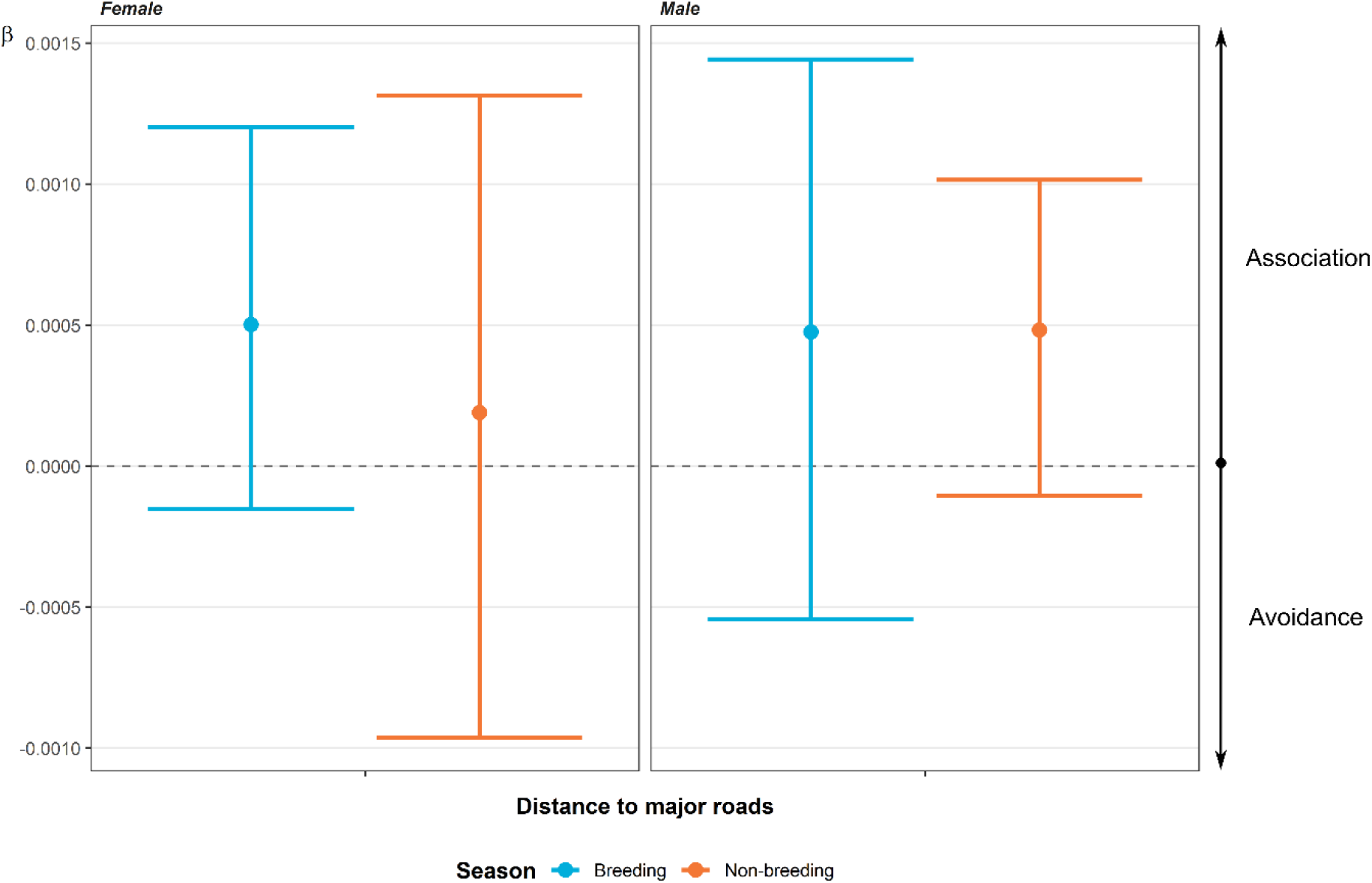
The coefficients relating to major roads from the population-level integrated step-selection function analyses. Breeding and non-breeding season are depicted by blue and orange respectively. Circles show the β estimate from the model and error bars show the associated 95% confidence intervals for each estimate.

## Discussion

We investigated the interactions of King Cobras with roads at the edge of a protected area within the Sakaerat Biosphere Reserve (SBR), Thailand. King Cobras repeatedly traversed a major four-lane highway, with individual snakes crossing up to 37 times during the study. We documented 32 potential crossing locations within a 15.3 km stretch of highway that telemetered King Cobras could potentially use to traverse the road. We observed three telemetered King Cobras using different types of underpasses, including one drainage culvert and two bridges (evidence of two; Figure 8). All individual-level ISSF results showed an equal or greater attraction to roads during the breeding season, except one individual, AM015, who we repeatedly observed moving in to the forest to breed and thus further away from road structures. Our population-level ISSF showed negligible changes in movement in relation to major roads for adult King Cobras within and outside the breeding season. However, our results suggest that adult female King Cobras exhibit a greater avoidance of roads during the non-breeding season, which we attribute to female King Cobras needing to cross a busy major road to access nesting sites during breeding season. Although all population-level ISSF confidence intervals overlapped zero, suggesting caution in interpreting our data, the evidence provided suggests that major roads bisecting typical female King Cobra occurrence distributions (within the agricultural landscape) and oviposition sites (forested areas) may present a particular mortality risk during what is already a hazardous time.

**Figure 8.**
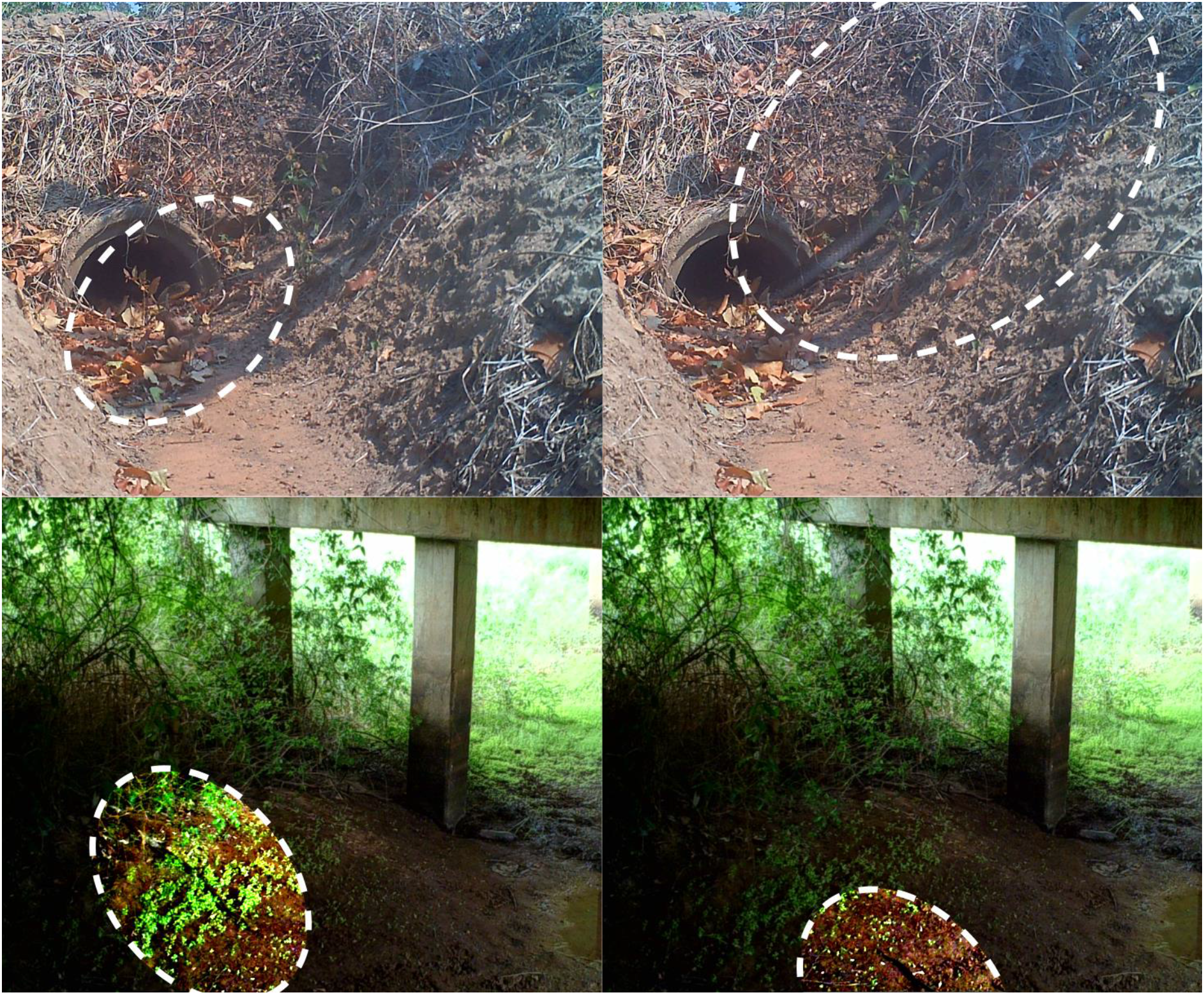
Use of road-crossing structures by telemetered King Cobra. *Top* Use of a drainage culvert by AM015. *Bottom* Movement underneath a bridge by AM054. King Cobras in frame are highlighted with dashed-white circles.

Intentional crossings structures for wildlife are absent from the SBR. We observed King Cobras in the SBR using a combination of drainage culverts and bridges to traverse the highway (Figure 4), but also attempting to cross the highway surface resulting in mortalities (Marshall et al., 2018). Because many of the telemetered King Cobras were tracked on one side of the road, and subsequently on the other side, we cannot directly confirm that individuals are routinely using underpasses as opposed to moving over the road. However, we have observed individuals moving underneath bridges (Figure 8), and we have obtained radio-telemetry fixes directly underneath Highway 304. Highway 304 is one of the busiest roads in Thailand; in 2006, 7,488 vehicles/day were recorded (Srikrajang 2006), which means one vehicle every 11.5 seconds on average. Moreover, with a consistent expansion of road infrastructure and users (Ng et al. 2020), we suspect traffic volumes to have been much higher during our study period than those observed in 2006. With all of our adult King Cobras having a total length between two and four metres, it is unlikely that an individual would be able to avoid a vehicle collision during a crossing event on the road surface. Given the contrast between these risks and the relatively high rate of successful crossings, we suspect that most King Cobras actively moving over the road are at high risk of vehicle collision, and are likely to be using under-the-road structures as presented in this study.

Adult males appeared to traverse the highway more frequently earlier in the year (February-May), which may be a result of mate searching behaviour during the breeding season (Marshall et al. 2019; Figure 3). Other snake species have shown a propensity to cross roads more frequently during breeding periods, due to increased mate searching activity (Bonnet et al. 1999; Lutterschmidt et al. 2005; Shepard et al. 2008b; Sosa and Schalk 2016). In our site, average crossing for adult males was much greater than for adult females, further supporting the mate searching role in road crossing frequency. However, this may reflect on the greater movement distance, frequency and overall greater occurrence distributions of adult males at our study site (Marshall et al. 2019).

Although we are confident that our unintentional crossing structures are being used by King Cobras to facilitate movement across the road, our low sampling frequency (∼1 pinpoint every 4-6 hours during the day) prevented us from routinely and confidently determining the structures used for every crossing (e.g., when multiple structures are available within close proximity of each other); we only have observations of King Cobras using three of our recorded structures. By inspecting our dBBMM subsets of crossing events, it is difficult to ascertain underpass use depending on individual, location and time of year. Supplementary Figure 4-7, for example, provide strong evidence of specific underpass use; however, Supplementary Figure 8-9, present ambiguous results due to individuals moving along the edge of the road prior to crossing. Additionally, we performed a single evaluation of each structure (haphazard temporal sampling) that did not allow us to detect seasonal changes in characteristics; for example, habitat connectivity and vegetation cover.

Our recursive analysis presented us with approximate dates and times that each road-crossing event occurred, and we subsequently investigated the points directly before and after a crossing in an attempt to discover if unintentional crossing structures were being used, and if so, which ones (Figure 4). Within the agricultural landscape of our study site, our results suggest that King Cobras used two bridges most frequently to cross the road (only one direct observation was made; Figure 8). The bridges are constructed over irrigation canals; the canals present an important landscape feature that appears to facilitate King Cobra movement throughout the agricultural landscape (Marshall et al. 2020). Research on other taxa suggests that the surrounding landscape provides the best predictor for which crossing structures are used over structural design and dimensions of culverts (Yanes et al. 1995; Rodríguez et al. 1996; Clevenger et al. 2002); however, we were unable to perform such analysis here due to the uncertainty surrounding if structures are actually being used. Therefore, it is difficult to determine whether King Cobras are selecting road crossing locations based on crossing structures presence (and the characteristics of the crossing structures), or if the landscape structure is funnelling individuals to these areas (i.e., connected to established movement corridors such as irrigation canals).

While encouraging, potential King Cobra use of road-crossing structures do not mitigate all Highway 304 potential impacts (Cunnington et al. 2014; Rytwinski et al. 2016). Throughout our study, we have encountered seven incidents of King Cobra road-mortality, five of which occurred on Highway 304. Out of these five highway-mortalities two were juvenile males, two were young of the year and one was a telemetered adult male. The newly hatched and juvenile snakes may be less acclimated to the presence of the crossing structures and distances between underpasses would be relatively greater and more challenging for smaller snakes to access; therefore, potentially increasing juvenile snakes’ vulnerability to road mortality. Several studies have reported increased road mortalities during juvenile emergence and dispersal (Erritzoe et al. 2003; Grilo et al. 2009; Kowalczyk et al. 2009). The discovery of our telemetered adult male, AM015, was worrying, particularly as our recursive analysis suggested that AM015 crossed underneath the same bridge on seven different occasions, showing a capacity to safely traverse the road, yet he crossed over the highway at least once and it led to the loss of the individual (Figure 4).

Reliance on underpasses may be insufficient to reduce road mortality, for small secretive taxa (such as amphibians) fencing and directive infrastructure are required to bolster underpasses’ effectiveness (Rytwinski et al., 2016). Rytwinski et al (2016) observations suggest that a combined approach of directive structures (e.g., fencing) and wildlife road crossings would better facilitate road crossing events for species that are at an increased risk of road mortality. However, King Cobras would require considerably more robust fencing than amphibians.

Female individuals of threatened taxa often require unique resources for reproduction (Brown and Weatherhead 1997; Roth and Greene 2006). Female King Cobras invest heavily in maternal care of eggs during oviposition and incubation (Whitaker et al. 2013; Hrima et al. 2014; Dolia 2018), and our long-term observations of King Cobra movement suggests that females shift their space-use during the nesting season to find suitable locations for oviposition (Marshall et al. 2019, 2020). In India, female King Cobras have been observed to remain with the nest post-laying (Whitaker et al. 2013; Hrima et al. 2014; Dolia 2018). Individual activity spikes (quantified by motion variance values from dynamic Brownian Bridge Movement Model output) were associated with King Cobra reproductive behaviours (i.e., oviposition site selection, nest guarding) at our site (Marshall et al. 2019). Female King Cobras in Thailand may travel into forested areas for nesting resources (e.g., substrate for nest building, vegetative cover and protection) unavailable in the agricultural matrix. Our results suggest that there are greater road mortality risks to reproductive female King Cobras during the pre- and post-nesting period (Marshall et al., 2020), when individual females ordinarily using agricultural areas make large, direct moves to forested areas in order to locate oviposition sites; typically putting these individuals at greater risk of encountering major roads.

Unintentional crossing structures (bridges and drainage culverts) appear to facilitate King Cobra movements across a fragmented landscape, providing some promise for the survival of the population in the presence of sizable human-made barriers like major roads. We have observed adult females moving beneath a bridge, allowing for safe passage across the 304NW-TLC road, but we have also observed females moving over the road surface, narrowly escaping oncoming vehicles. Allocation of designed wildlife crossing structures along both of our sampled roads, using guidance from our movement data and previously outlined mortality hotspots in Silva et al. (2020), could provide a foundation for plans to reduce mortality caused by these roads for a diversity of taxa within the Sakaerat Biosphere Reserve.

## Conclusion

Our findings add to a growing collection of road ecology literature attempting to decipher how animals interact with anthropogenic obstacles. Unintentional ecological underpasses are likely providing some level of permeability and prevent complete habitat fragmentation, which is particularly important for snakes given their reluctance to cross roads and their vulnerability when doing so (Shine et al. 2004; Andrews and Gibbons, 2005). King Cobras being larger and ranging further than most other reptiles makes them additionally vulnerable to habitat fragmentation and the dangers of roads (Bonnet et al. 1999; Rytwinski and Fahrig 2012). The presence of crossing structures (drainage culverts and bridges) along a major four-lane highway appears to enable King Cobras to traverse the road, providing a level of permeability. Despite this, we continue to discover individuals that have died due to vehicle collision on Highway 304. We suggest two main future study avenues to be explored at the Sakaerat Biosphere Reserve. First, a monitoring study should be designed to evaluate the true use of unintentional wildlife crossing structures, as presented in this study, either via strategic and coordinated camera traps, or via more advanced systems, such as PIT-tag readers (Bateman et al. 2017). Second, research is needed to evaluate whether guidance fencing combined with a Before After Control Impact (BACI) along the Highway 304 structures could aid in limiting road mortalities for both King Cobras and other terrestrial species.

## Acknowledgements

We thank the National Park, Wildlife and Plant Conservation Department, Thailand for giving us permission to study a nationally protected species, and the National Research Council of Thailand for providing permits to undertake this research. With these permissions, we are grateful for surgical expertise provided by Nakhon Ratchasima Zoo, Dusit Zoo, and Zoological Park Organization under the Royal Patronage of His Majesty the King, Thailand.

We thank the Institute of Animal Scientific Purpose Development for providing C.T.S. and P.S. with animal use licenses. We thank the Suranaree University of Technology and the School of Biology for ethically approving the study and further providing logistical, supervisory and financial aid. We thank Pluemjit Boonpueng for assisting with paper work and logistics. We thank Wildlife Reserves Singapore Conservation Fund, National Scientific and Technological Development Agency and Thailand Institute of Scientific and Technological Research, and the National Research Council of Thailand for funding the research at different stages. We also thank the Herpetofauna Foundation and St Stephen’s International School, Bangkok, for donating essential equipment. We thank the staff and management at Sakaerat Environmental Research Station for their long-term logistic support throughout the project. We thank the residents of Udom Sab for allowing research to be undertaken across their land. We thank the Hook 31 Rescue teams for their tireless work mitigating human-snake conflict and providing us with a number of King Cobras. We give special thanks to the many Sakaerat Conservation and Snake Education Team members for dedicating countless hours tracking King Cobras throughout the landscape.

## Availability of data and materials

Data used in this study is available on Zenodo (DOI: 10.5281/zenodo.5148436) and Movebank (Movebank ID: 1649411628). The Zenodo repository also includes all R scripts used to run analysis.

## Author Contributions

*Conceptualization*, M.D.J., B.M.M., S.N.S. and C.T.S.; *Methodology*, M.D.J., B.M.M, S.N.S. and C.T.S; *Formal Analysis*, M.D.J., B.M.M and S.N.S.; *Investigation*, M.D.J., B.M.M., S.N.S., M.C., I.S. and C.T.S., *Writing – Original Draft*, M.D.J.; *Writing – Review & Editing*, M.D.J., S.N.S., C.T.S., B.M.M., M.C., I.S., W.W., M.G, P.S., T.A., S.W.; *Visualization*, M.D.J. and B.M.M.; *Supervision*, P.S., S.W., and T.A.; *Funding Acquisition*, M.D.J., C.T.S., and P.S.

## Ethics approval and consent to participate

We had ethical approval from the Suranaree University Ethics Committee (24/2560). Our study was undertaken using the Institute of Animals for Scientific Purpose Development (IAD) licenses belonging to P.S and C.T.S. Permission was granted by the National Park, Wildlife and Plant Conservation Department and Thailand and the National Research Council of Thailand (98/59). Further permission for work was given by the Thailand Institute of Scientific and Technological Research and Sakaerat Environmental Research Station.

