## Supplementary File for "How do King Cobras move across a major highway? Unintentional wildlife crossing structures may facilitate movement"

**Affiliations**:

We completed all analyses in R v.3.5.3 (R Core Team 2019) and R Studio v.1.2.1335 (R Studio Team 2019). We performed all data manipulation using R packages *dplyr* v.0.8.3 (Wickham et al. 2019), *lubridate* v.1.7.4 (Grolemund and Wickham 2011), *readr* v.1.3.1 (Wickham and Hester 2020), *reshape2* v.1.4.3 (Wickham 2007), and *stringr* v.1.4.0 (Wickham 2019). We calculated data means and standard error (±) using the *pracma* package v.2.2.5 (Borchers 2019). We worked with rasters and shapefiles (for producing study site maps and recurse analysis) using R packages *raster* v.2.8.19 (Hijmans 2019), *rgdal* v.1.4.3 (Bivand et al. 2019) and *sp* v.1.3.1 (Pebesma and Bivand 2005; Bivand et al. 2013). We created visuals using a combination of R packages *cowplot* v.0.9.4 (Wilke 2019), *ggplot2* v.3.2.1 (Wickham 2009), *ggspatial* v.1.0.3 (Dunnington 2018), *scales* v.1.1.0 (Wickham and Seidel 2019) and *scico* v.1.1.0 (Pederson and Crameri 2018).

Supplementary Table 1. Capture and release information for all telemetered King Cobra.

| ID | Capture Date | Capture Easting | Capture Northing | Release Date | Release Easting | Release Northing | Distance (m) |
|---|---|---|---|---|---|---|---|
| AM006 | 5th April 2018 | 818746 | 1605879 | 10th April 2018 | 819064 | 1605795 | 329 |
| AM007 | 30th March 2019 | 818552 | 1604731 | 2nd April 2019 | 818552 | 1604731 | 0 |
| AF010* | 6th March 2015 | 818918 | 1607399 | 15th March 2017 | 818926 | 1607392 | 11 |
| JM013* | 6th July 2014 | 818471 | 1606679 | 19th July 2014 | 818267 | 1606533 | 251 |
| AM015* | 11th October 2014 | 818222 | 1606078 | 26th October 2014 | 817935 | 1606003 | 297 |
| AF017* | 28th April 2015 | 818424 | 1607178 | 6th May 2015 | 818376 | 1607172 | 48 |
| AM018* | 28th April 2015 | 818002 | 1605882 | 9th May 2015 | 817789 | 1605758 | 246 |
| JM019* | 1st November 2015 | 820547 | 1605330 | 7th November 2015 | 820842 | 1604127 | 1263 |
| AM024 | 25th April 2016 | 815815 | 1605104 | 1st May 2016 | 815859 | 1605160 | 71 |
| JM025 | 25th May 2016 | 817145 | 1606125 | 31st May 2016 | 817145 | 1606125 | 0 |
| AM026 | 28th November 2016 | 814706 | 1602980 | 30th November 2016 | 814706 | 1602980 | 0 |
| JF027* | 14th January 2017 | 818809 | 1606327 | 15th January 2017 | 818968 | 1606376 | 166 |
| JM034* | 24th April 2017 | 820227 | 1606327 | 18th May 2017 | 820351 | 1605016 | 473 |
| AM054 | 28th February 2018 | 818726 | 1609537 | 2nd March 2018 | 818933 | 1608996 | 582 |
| JF055 | 14th March 2018 | 812944 | 1600279 | 16th March 2018 | 812993 | 1600065 | 221 |
| AF056 | 24th March 2018 | 817836 | 1608698 | 29th March 2018 | 817836 | 1608698 | 0 |
| AF058 | 6th April 2018 | 816730 | 1604030 | 10th April 2018 | 816730 | 1604030 | 0 |
| AM059 | 28th March 2019 | 820302 | 1607863 | 2nd April 2019 | 820241 | 1607857 | 61 |
| AF086 | 23rd April 2019 | 820274 | 1608801 | 25th April 2019 | 820274 | 1608801 | 0 |
| AF096 | 10th March 2020 | 821702 | 1610664 | 16th March 2020 | 821702 | 1610664 | 0 |
| AF099 | 18th March 2020 | 820608 | 1607174 | 23rd March 2020 | 820608 | 1607174 | 0 |

ID* depict individuals where true release sites were not recorded and the first datapoint was used. *Distance* Distance between capture and release UTM’s.

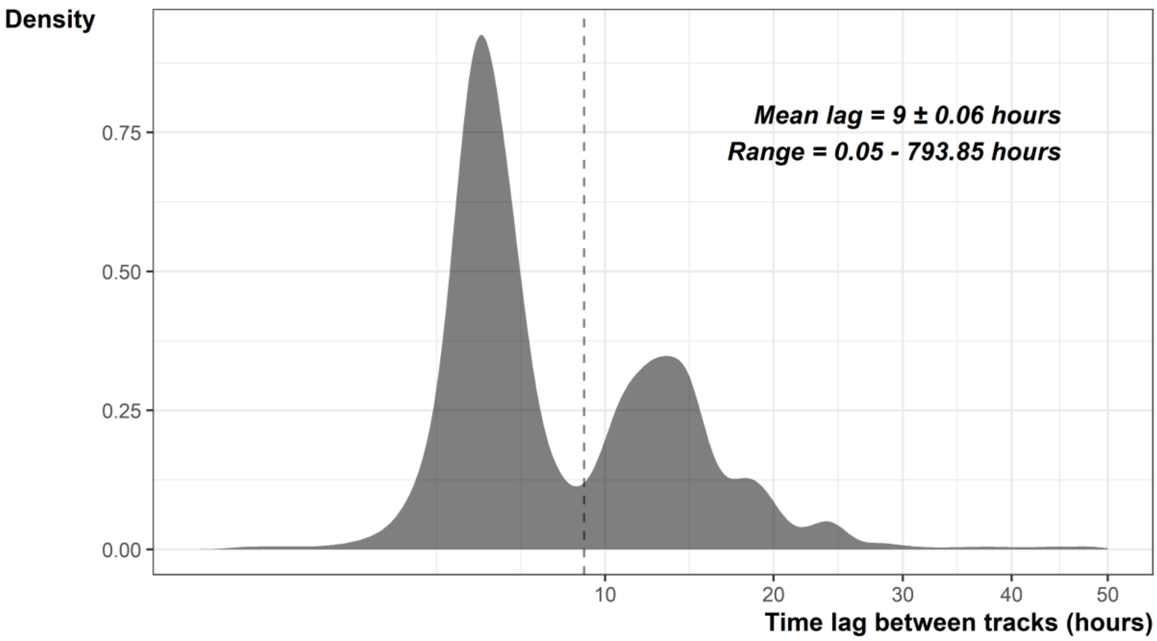


Supplementary Figure 1. Distribution of time lags between datapoints. Horizontal line depicts the mean time lag.

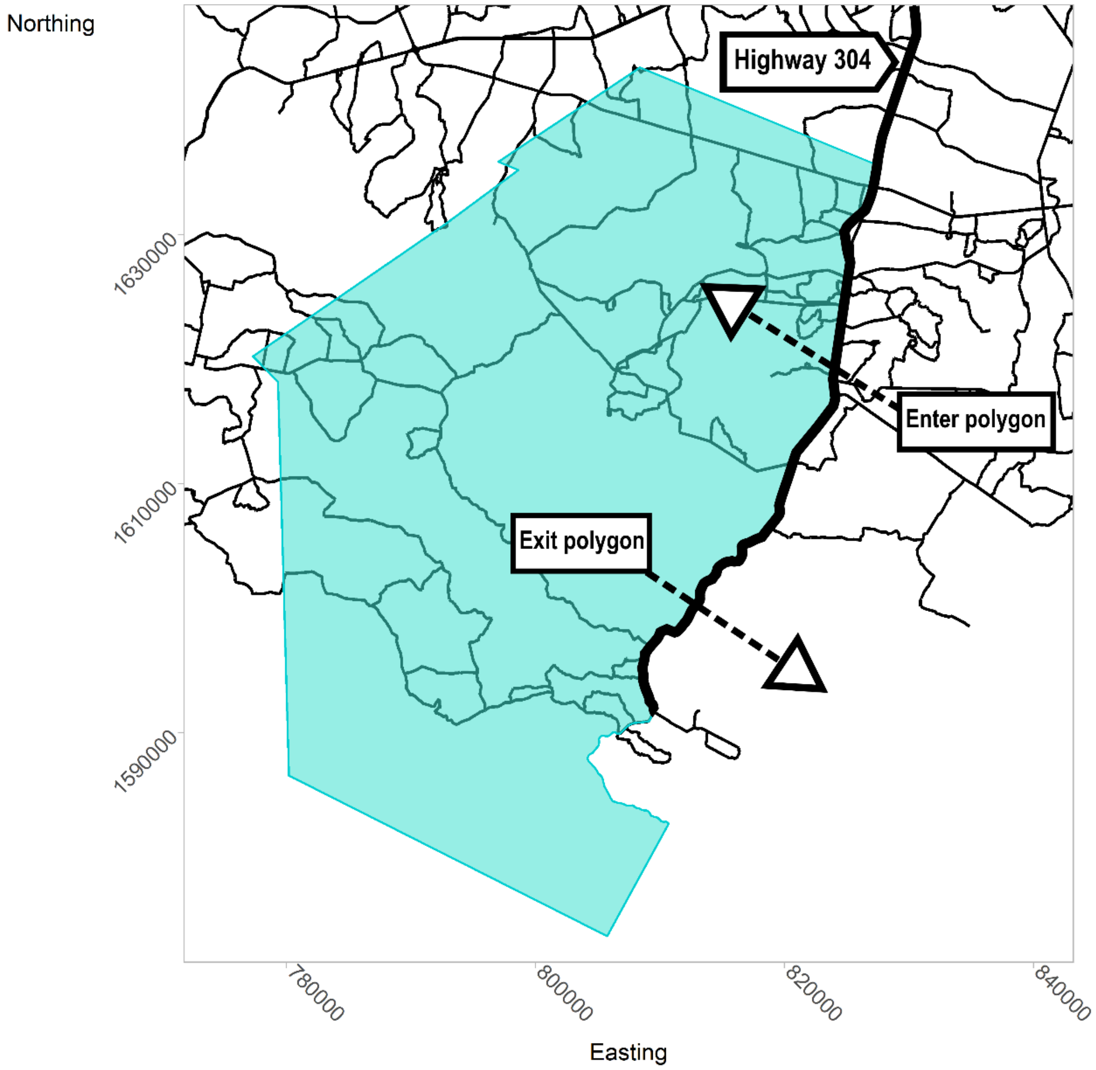


Supplementary Figure 2. The North Side polygon (blue) used in the recurse analysis to determine road-crossings across the Highway 304.

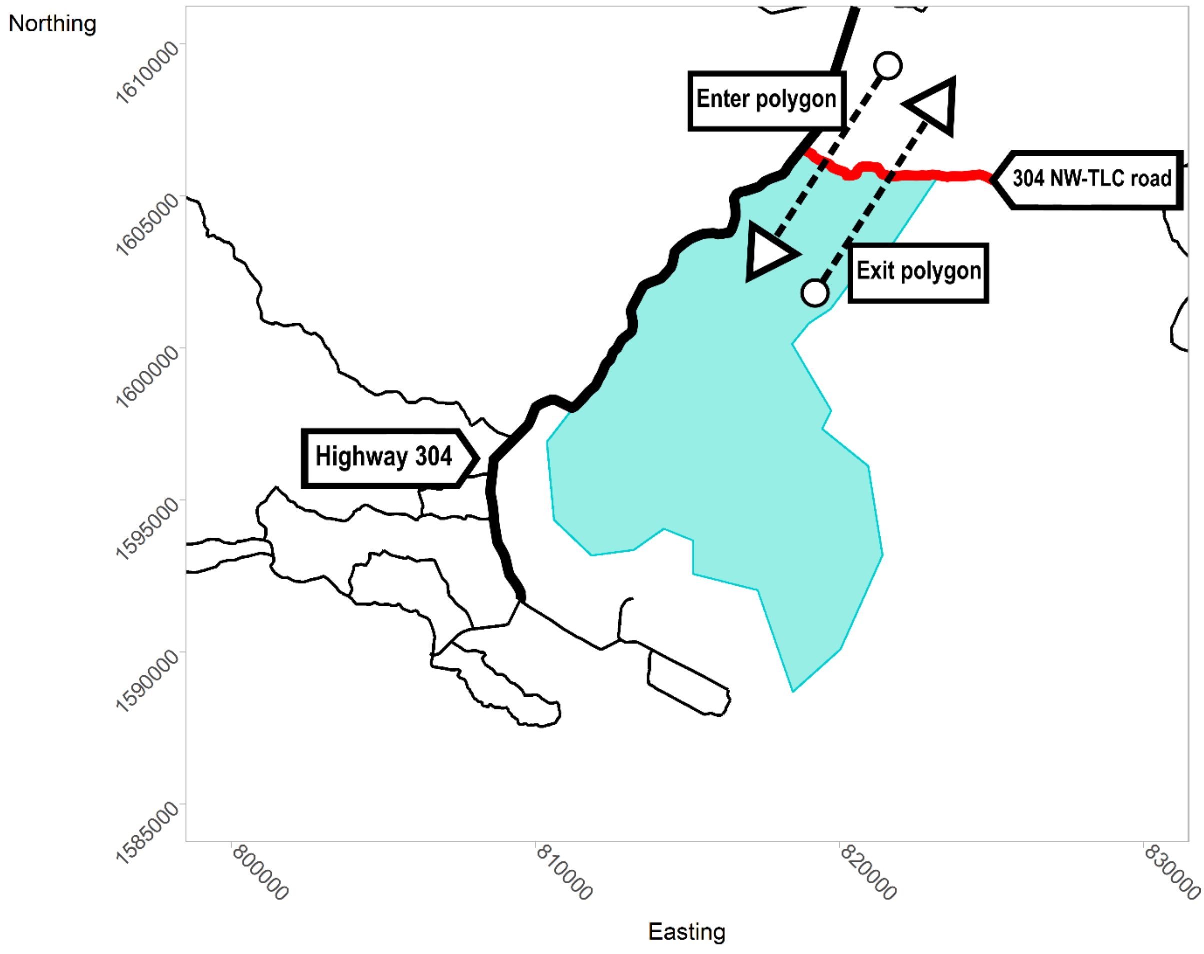


Supplementary Figure 3. The South Side polygon (blue) used in the recurse analysis to determine road-crossings across the 304 NW-TLC road.

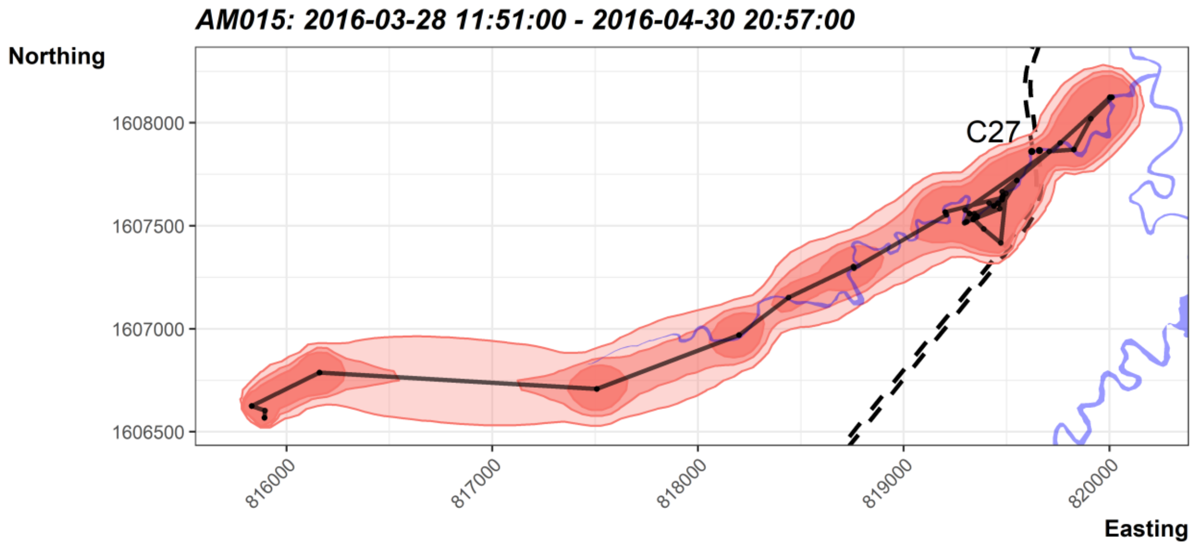


Supplementary Figure 4. 90, 95 and 99% use contours estimated using dynamic Brownian Bridge Movement Models of AM015 during a road-crossing event. 90, 95 and 99% contours are differentiated by increasing opacity. Solid black lines represent the movement trajectory of the King Cobra. Highway 304 is depicted using black dashed lines and crossing structures are labelled next to their position on Highway 304. Blue polygons show the irrigation canals. Map is in a North up orientation.

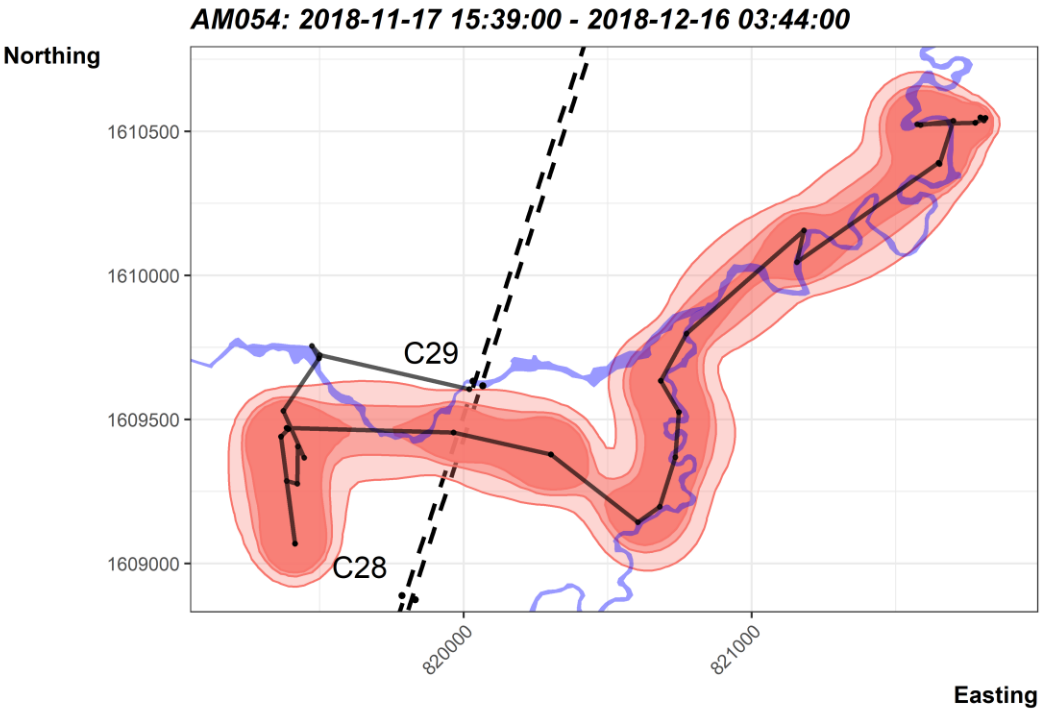


Supplementary Figure 5. 90, 95 and 99% use contours estimated using dynamic Brownian Bridge Movement Models of AM054 during a road-crossing event. 90, 95 and 99% contours are differentiated by increasing opacity. Solid black lines represent the movement trajectory of the King Cobra. Highway 304 is depicted using black dashed lines and crossing structures are labelled next to their position on Highway 304. Blue polygons show the irrigation canals. Map is in a North up orientation.

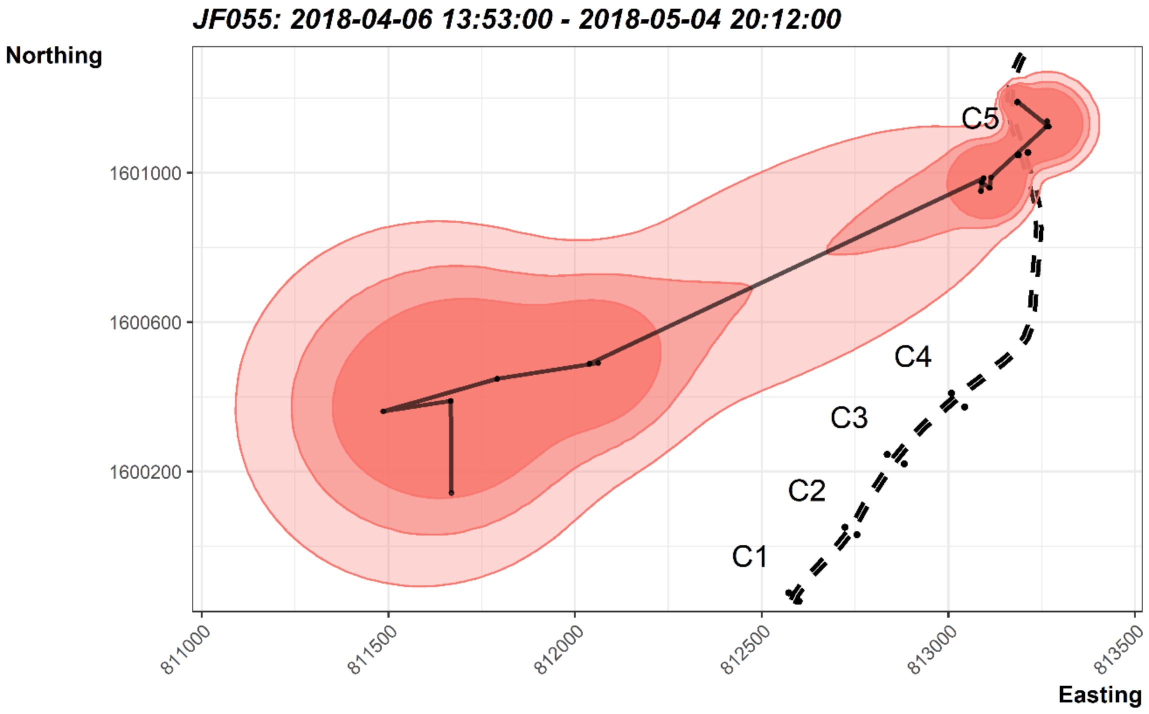


Supplementary Figure 6. 90, 95 and 99% use contours estimated using dynamic Brownian Bridge Movement Models of JF055 during a road-crossing event. 90, 95 and 99% contours are differentiated by increasing opacity. Solid black lines represent the movement trajectory of the King Cobra. Highway 304 is depicted using black dashed lines and crossing structures are labelled next to their position on Highway 304. Blue polygons show the irrigation canals. Map is in a North up orientation.

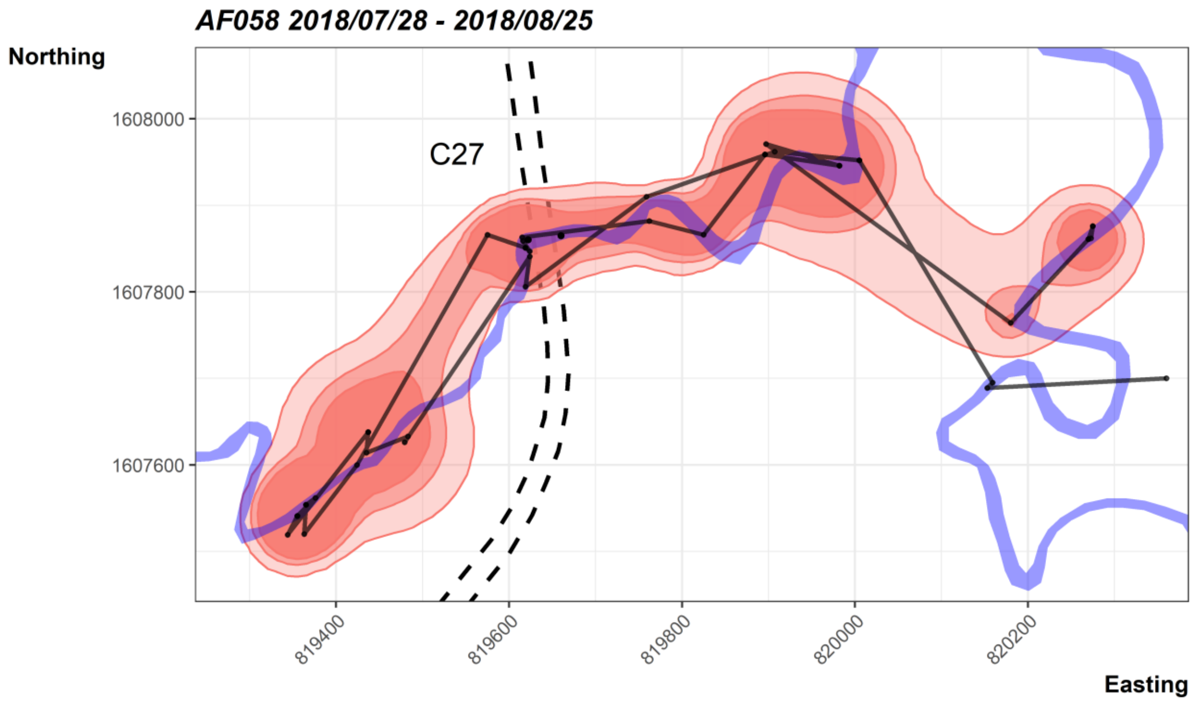


Supplementary Figure 7. 90, 95 and 99% use contours estimated using dynamic Brownian Bridge Movement Models of AF058 during a road-crossing event. 90, 95 and 99% contours are differentiated by increasing opacity. Solid black lines represent the movement trajectory of the King Cobra. Highway 304 is depicted using black dashed lines and crossing structures are labelled next to their position on Highway 304. Blue polygons show the irrigation canals. Map is in a North up orientation.

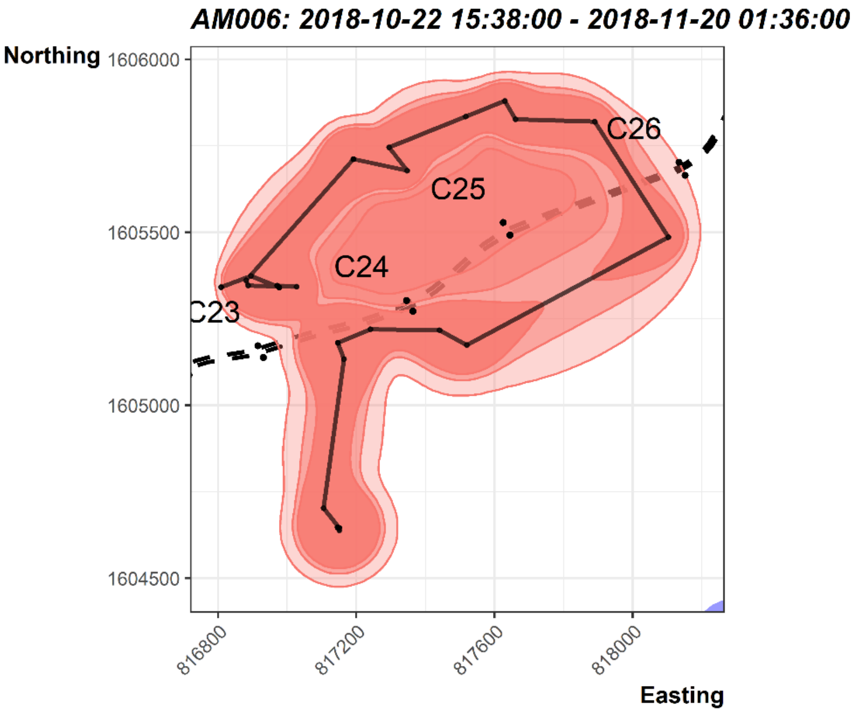


Supplementary Figure 8. 90, 95 and 99% use contours estimated using dynamic Brownian Bridge Movement Models of AM006 during a road-crossing event. 90, 95 and 99% contours are differentiated by increasing opacity. Solid black lines represent the movement trajectory of the King Cobra. Highway 304 is depicted using black dashed lines and crossing structures are labelled next to their position on Highway 304. Blue polygons show the irrigation canals. Map is in a North up orientation.

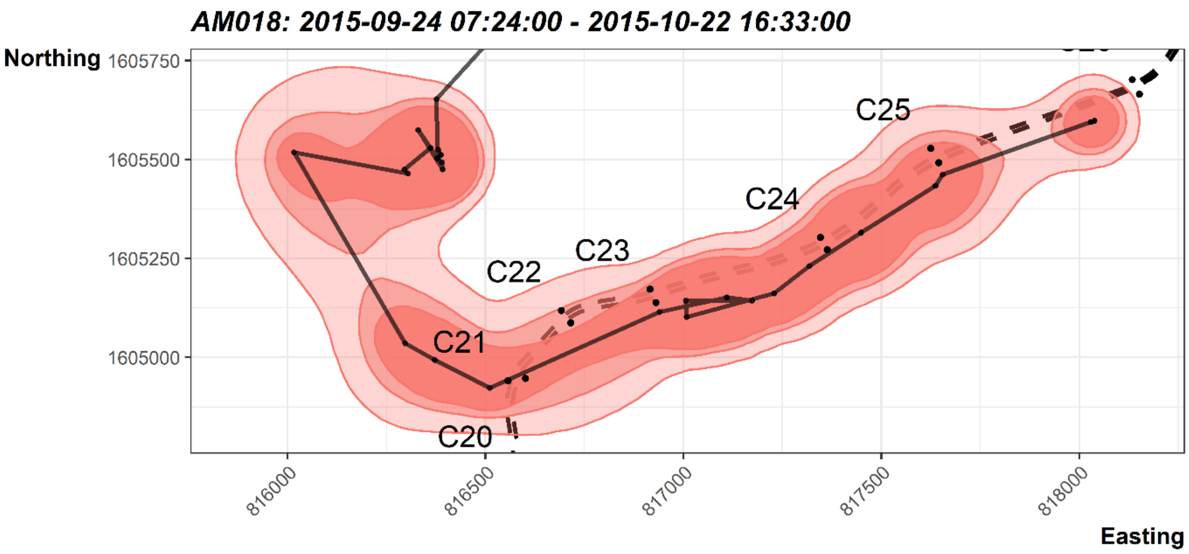


Supplementary Figure 9. 90, 95 and 99% use contours estimated using dynamic Brownian Bridge Movement Models of AM018 during a road-crossing event. 90, 95 and 99% contours are differentiated by increasing opacity. Solid black lines represent the movement trajectory of the King Cobra. Highway 304 is depicted using black dashed lines and crossing structures are labelled next to their position on Highway 304. Blue polygons show the irrigation canals. Map is in a North up orientation.

Supplementary Table 2. Characteristics of road crossing structures. *Type* one of three structure types: culvert, double culvert or bridge. *Times used* the number of times the structure was used to traverse the Highway 304 by telemetered King Cobra. *Length* straight line distance of the structure from entrance one to entrance two. *Width* the horizontal measurement (mm) of the structure entrance. *Height* the vertical measurement (mm) of the structure entrance. *Substrate* the dominant substrate type present within the structure. *Material* the dominant material types the structure is comprised of. *Vegetation* whether or not there was vegetative cover at the entrance of the structure. *Waste* whether there was anthropogenic waste immediately outside, or within, the structure. *Feature* the aquatic landscape feature that the structure led into. *Distance* mean Euclidean distance to the two nearest crossing structures.

| | Type | Times used | Entrance one | | Entrance two | | Length | Width | Height | Substrate | Material | Vegetation | Waste | Feature | Distance |
|---|---|---|---|---|---|---|---|---|---|---|---|---|---|---|---|
| | | | Easting | Northing | Easting | Northing | | | | | | | | | |
| C1 | Culvert | 1 | 812573 | 1599875 | 812600 | 1599853 | 35 | 942 | 851 | None | Concrete | Yes | Yes | None | 230 |
| C2 | Culvert | 0 | 812723 | 1600051 | 812755 | 1600031 | 37 | 981 | 986 | None | Concrete | Yes | Yes | None | 227 |
| C3 | Culvert | 1 | 812836 | 1600246 | 812882 | 1600221 | 53 | 983 | 995 | None | Concrete | Yes | Yes | Stream | 231 |
| C4 | Culvert | 0 | 813008 | 1600410 | 813044 | 1600373 | 52 | 975 | 984 | None | Concrete | No | Yes | None | 449 |
| C5 | Bridge | 2 | 813187 | 1601048 | 813213 | 1601054 | 26 | 22000 | 2422 | Gravel | Concrete | Yes | Yes | Klong | 623 |
| C6 | Culvert | 0 | 813353 | 1601611 | 813392 | 1601596 | 42 | 962 | 992 | None | Concrete | No | Yes | None | 459 |
| C7 | Culvert | 1 | 813495 | 1601913 | 813527 | 1601895 | 37 | 898 | 874 | None | Concrete | Yes | Yes | None | 322 |
| C8 | Culvert | 1 | 813724 | 1602127 | 813745 | 1602091 | 41 | 943 | 782 | None | Concrete | Yes | Yes | None | 338 |
| C9 | Culvert | 0 | 814065 | 1602257 | 814082 | 1602218 | 42 | 966 | 667 | Rocks | Concrete | Yes | Yes | None | 324 |
| C10 | Culvert | 1 | 814449 | 1602612 | 814475 | 1602591 | 33 | 975 | 995 | None | Concrete | Yes | Yes | None | 264 |
| C11 | Culvert | 0 | 814296 | 1602424 | 814322 | 1602403 | 33 | 514 | 194 | None | Concrete | Yes | Yes | None | 386 |
| C12 | Culvert | 0 | 814587 | 1603127 | 814665 | 1603101 | 82 | 930 | 597 | None | Concrete | Yes | Yes | Stream | 397 |
| C13 | Double Culvert | 2 | 814769 | 1603324 | 814795 | 1603291 | 42 | 906 | 643 | Gravel | Concrete | Yes | Yes | Stream | 303 |
| C14 | Culvert | 1 | 815033 | 1603534 | 815052 | 1603507 | 34 | 2100 | 1790 | None | Concrete | No | Yes | Stream | 316 |
| C15 | Double Culvert | 1 | 815298 | 1603666 | 815315 | 1603636 | 35 | 975 | 960 | Rocks | Concrete | Yes | Yes | Stream | 263 |
| C16 | Culvert | 2 | 815515 | 1603745 | 815519 | 1603707 | 39 | 990 | 985 | None | Concrete | Yes | Yes | Stream | 212 |
| C17 | Culvert | 0 | 815706 | 1603768 | 815710 | 1603736 | 33 | 982 | 984 | None | Concrete | Yes | Yes | None | 444 |
| C18 | Double Culvert | 3 | 816402 | 1603809 | 816425 | 1603780 | 38 | 897 | 967 | None | Concrete | Yes | Yes | None | 630 |
| C19 | Culvert | 1 | 816602 | 1604334 | 816634 | 1604330 | 32 | 970 | 993 | None | Concrete | Yes | Yes | None | 464 |
| C20 | Culvert | 3 | 816569 | 1604701 | 816604 | 1604704 | 35 | 950 | 979 | Rocks | Concrete | Yes | Yes | None | 607 |
| C21 | Double Culvert | 14 | 816557 | 1604941 | 816601 | 1604947 | 45 | 1040 | 951 | Gravel | Concrete | Yes | Yes | Stream | 231 |
| C22 | Culvert | 3 | 816692 | 1605118 | 816716 | 1605087 | 39 | 960 | 942 | Waste | Concrete | Yes | Yes | Stream | 225 |
| C23 | Culvert | 1 | 816916 | 1605172 | 816931 | 1605138 | 37 | 972 | 930 | None | Concrete | Yes | Yes | None | 338 |
| C24 | Double Culvert | 3 | 817346 | 1605303 | 817364 | 1605272 | 35 | 925 | 935 | None | Metal | Yes | No | Stream | 402 |

| | | | | | | | | | | | | | | | |
|---|---|---|---|---|---|---|---|---|---|---|---|---|---|---|---|
| C25 | Bridge | 1 | 817625 | 1605528 | 817645 | 1605492 | 40 | 12000 | 1750 | Rocks | Concrete | Yes | Yes | None | 447 |
| C26 | Double Culvert | 20 | 818134 | 1605702 | 818152 | 1605665 | 41 | 965 | 983 | Soil | Concrete | Yes | Yes | None | 1579 |
| C27 | Bridge | 18 | 819622 | 1607860 | 819660 | 1607865 | 38 | 30000 | 3830 | Soil | Concrete | Yes | Yes | Klong | 1827 |
| C28 | Culvert | 0 | 819786 | 1608888 | 819832 | 1608874 | 49 | 800 | 480 | Water | Concrete | No | Yes | None | 910 |
| C29 | Bridge | 6 | 820033 | 1609633 | 820067 | 1609617 | 37 | 30000 | 3000 | Gravel | Concrete | Yes | Yes | Klong | 2131 |
| C30 | Culvert | 0 | 820446 | 1610913 | 820491 | 1610896 | 49 | 850 | 980 | Water | Concrete | Yes | Yes | None | 1321 |
| C31 | Culvert | 0 | 820848 | 1612152 | 820896 | 1612135 | 51 | 1000 | 1000 | Gravel | Concrete | Yes | Yes | None | 881 |
| C32 | Double Culvert | 0 | 821022 | 1612582 | 821062 | 1612557 | 48 | 1000 | 1000 | Water | Concrete | Yes | Yes | None | 465 |
